# Acoustic array biochip combined with allele-specific PCR for multiple cancer mutation analysis in tissue and liquid biopsy

**DOI:** 10.1101/2021.09.16.460590

**Authors:** Nikoletta Naoumi, Kleita Michaelidou, George Papadakis, Agapi E. Simaiaki, Román Fernández, Maria Calero, Antonio Arnau, Achilleas Tsortos, Sofia Agelaki, Electra Gizeli

## Abstract

Regular screening of cancerous point mutations is of importance to cancer management and treatment selection. Although excellent techniques like next generation sequencing and droplet digital PCR are available, these are still lacking in speed, simplicity and cost-effectiveness. Here a new approach is presented where allele-specific PCR (AS-PCR) is combined with a novel High Fundamental Frequency Quartz Crystal Microbalance (HFF-QCM) array biosensor for the amplification and detection, respectively, of cancer point mutations. For the proof-of-concept, the method was applied to the screening of the *BRAF* V600E and *KRAS* G12D mutations in spiked-in and clinical samples. Regarding the BRAF target, an analytical sensitivity of 0.01%, i.e., detection of 1 mutant copy of genomic DNA in an excess of 10^4^ wild type molecules, was demonstrated; moreover, quantitative results during KRAS detection were obtained when an optimized assay was employed with a sensitivity of 0.05%. The assays were validated using tissue and plasma samples obtained from melanoma, colorectal and lung cancer patients. Results are in full agreement with Sanger sequencing and droplet digital PCR, demonstrating efficient detection of *BRAF* and *KRAS* mutations in samples having an allele frequency below 1%. The high sensitivity and technology-readiness level of the methodology, together with the ability for multiple sample analysis (24 array biochip), cost-effectiveness and compatibility with routine work-flow, hold promise for the implementation of this AS-PCR/acoustic methodology in clinical oncology as a tool for tissue and liquid biopsy.

## 1. Introduction

Tumor tissue biopsy (TB) remains the gold standard method for cancer diagnosis and is an important source for routine molecular profiling of hotspot somatic mutations. However, conventional tissue biopsy still has a number of limitations that stem from its invasive nature. In many cases, is not feasible to perform TB and is inappropriate for capturing intra-tumor heterogeneity, longitudinal profiling of cancer biomarkers and monitoring of disease progression^1,2,3^.

Liquid biopsy (LB) is a promising non-invasive alternative strategy to tissue biopsy that allows the study and characterization of different biomarkers such as cell-free DNA fragments originating from tumor cells. Circulating tumor DNA (ctDNA) may carry the same genetic alterations as those of a primary tumor and thus, can serve as a valuable diagnostic and prognostic tool to select targeted therapies and monitor therapeutic response in real time^4^. While LB is simpler, faster and more cost-effective than TB, ctDNA detection poses an analytical challenge since it is present in very small quantities (z/aM), is highly fragmented and exists in samples with a background of abundant wild type (wt) cell free DNA, exhibiting a ratio of mt:wt below 0.1 %^5^.

Generally, mutation analysis of ctDNA can be performed by two different approaches: next generation sequencing (NGS) and PCR-based methods^6,7^. Although NGS has contributed considerably to molecular diagnostics for clinical oncology, its application is restricted by the high cost, complexity and slow turnaround time (up to 10 days); in addition, only a small number of labs are equipped with a NGS facility. Real time PCR (qPCR) is also applied routinely for cancer molecular analysis. The limitations of qPCR, compared to NGS-based methods, include the low sensitivity of detecting mutations in the presence of wild type DNA (Mutant allele frequency; MAF of 10-20%)^4^, and the need for targeted mutation analysis^8,9^. The Allele-Specific PCR (AS-PCR) and the third-generation droplet digital-PCR (ddPCR) are able to detect 0.1% and 0.001% MAFs, respectively^7^. ddPCR has limitations including long droplet processing times and high cost while AS-PCR, although more cost-effective, exhibits low sensitivity during the amplification of cancer mutant (mt) targets in the presence of an excess of background wild-type (wt) cell-free DNA.

Recently, DNA biosensors, including electrochemical^10–12^, optical^13,14^ and acoustic^15–18^, have been reported as an alternative means for the detection of point mutations. Moreover, advances in biochips and nanotechnology have led to the development of enzymatic amplification-free protocols for ctDNA^7,19^. Such examples include an electrochemical biochip combined with surface nanostructuring and a clutch/clamp assay^20^; a plasmonic biosensor employing nanoparticles and a PNA-functionalized gold surface^21^; and a piezoelectric plate sensor combined with fluorescent reporter microspheres^22^. Overall, the above works present elegant examples of the application of biosensors in PCR-free clinical diagnostics, often achieving impressive sensitivities similar to that of ddPCR^20^. Potential drawbacks include the (a) use for expensive peptide nucleic acid (PNA) or locked nucleic acid (LNA) probes for the specific capturing of the mt target; (b) requirement for hybridization steps and temperature control; (c) laborious surface-activation and washing steps as well as nanoparticles synthesis and functionalization; and (d) low sensitivity, in cases when low volumes of unpurified sample is used.

Because during the past decade LB has received tremendous attention, there is an urgent need for the development of diagnostic tools for immediate application in the clinic^23^. Such tools should, ideally, be rapid and simple to use; compatible to the routine methods currently employed in diagnostic labs for fast adaptation; exhibit improved sensitivity and specificity; allow for multi-analysis; be robust; and, finally provide a more economical solution to current methods. Inspired by the above needs, we developed a method for the detection of point-mutations utilizing a novel High Fundamental Frequency Quartz Crystal Microbalance (HFF-QCM) array-device with dissipation monitoring allowing multi-sample analysis through the use of 24 resonators. Acoustic detection is coupled with AS-PCR for the amplification of 1-10^3^ mt DNA targets carrying the *BRAF* V600E or *KRAS* G12D point mutation in a background of 10^4^ wt molecules. To combine AS-PCR with acoustic detection, we designed the assay so that the double strand DNA (dsDNA) products could be directly immobilized on the device surface, obviating the need for a hybridization step. Furthermore, we employed liposomes as acoustic signal enhancers in order to improve the detection capability of the assay and detect down to 1 mutant copy in the initial sample in the presence of 10^4^ wild type DNAs.

The clinical validity of the assay was further demonstrated during the successful detection of *BRAF* and *KRAS* point mutations in colorectal, lung and melanoma cancer patients’ tissue and plasma samples. Results indicate excellent sensitivity of the method, comparable to that obtained with ddPCR, with a more simple and less expensive manner.

## 2. Materials and methods

### 2.1 Acoustic array and QCMsensor

The 150 MHz HFF-QCM acoustic array signals were monitored with a newly developed platform (AWS, S.L. Paterna, Spain). The QCM sensors (AWS, S.L. Paterna, Spain), with a fundamental frequency of 5 MHz, were monitored using the Q-Sense E4 instrument (Biolin Scientific, Sweden) at the 7th overtone (35MHz). Details on the device cleaning can be found in the Supplementary S1.

### 2.2 Acoustic detection of b-BSA and NAv

Protein samples were diluted in PBS pH=7.4 (Sigma-Aldrich). Neutravidin (NAv-Invitrogen) (0.2 mg/mL) and biotinylated-BSA (b-BSA) (0.2 mg/mL) were applied on the device surface; 0.05 mg/mL of NAv was further applied on the b-BSA layer. b-BSA was prepared as described in SI-S2. Due to the different size of the HFF QCM and QCM devices and fluidics geometry, the working volumes were V_150MHz_=60 μL and V_35MHz_=200 μL.

### 2.3 HFF QCM detection of dsDNA and liposomes

dsDNA fragments of 21 bp, 50 bp 75 bp and 157 bp were prepared according to^24^ and applied (60 μL of 83 or 500 nM) to a NAv pre-coated array. 100 μL x 0.2 mg/mL of 200 nm of POPC liposomes (1-palmitoyl-2-oleoyl-glycero-3-phosphocholine), prepared by extrusion as described before^24^, were added on the DNA surface.

### 2.4 Acoustic analysis of AS-PCR

For the acoustic detection of amplicons, the gold sensor surface was modified with b-BSA followed by the addition of NAv (see above). A sample of 2.5 μL or 8 μL of the *BRAF* or *KRAS* AS-PCR reaction respectively, diluted in a total volume of 20 μL, was loaded on the 150 MHz HFF QCM array (flow rate: 14 μL/min). Similarly, 2.5 μL or 10 μL of the *BRAF* or *KRAS* AS-PCR, diluted in a total volume of 125 μL, was applied to the 35 MHz QCM device at a flow rate of 25 μL/min. In both cases, a suspension of 0.2 mg/mL of 200 nm POPC liposomes was added at a volume of 100 μL (150 MHz) or 500 μL (35 MHz).

### 2.5 BRAF V600E and KRAS G12D AS-PCR

For the *BRAF* V600E, KAPA2G Fast HotStart ReadyMix (KAPABIOSYSREMS) was mixed with 5 pmol of the allele-specific biotinylated reverse primer and with 5 pmol of the cholesterol modified forward primer in a total volume of 10 μL. For the *KRAS* G12D, 10 pmol of the mutation-specific biotinylated forward primer, 10 pmol of the cholesterol-modified reverse primer and 1μL 20X SYBR^®^ Green I Nucleic Acid Stain (Lonza) were mixed with the KAPA2G Fast HotStart ReadyMix in a total volume of 20 μL. More information is provided in the SI-S3 and Table S1.

### 2.6 Sample collection

21 FFPE tumor and 20 plasma samples were obtained from patients with various cancer types at the University Hospital of Heraklion. The research protocol was approved by the Institutional Ethics Committee of the University Hospital and all patients provided written informed consent to participate in the study.

### 2.7 Sanger Sequencing and ddPCR for KRAS and BRAF-mutation analysis

gDNA from FFPE tissues, was amplified by PCR using specific primer pairs for *KRAS* exon 2 and *BRAF* exon 15 (Table S1). Sequencing reactions were performed using the Big Dye terminator V3.1 cycle sequencing kit (Applied Biosystems) according to the manufacturer’s protocol. The products were then assessed by capillary electrophoresis on an ABI3130 System and results were analyzed using Sequencing Analysis software v5.4 (Applied Biosystems).

cfDNA was isolated from 2 mL of plasma for each sample via the QIAamp Circulating Nucleic Acid kit (Qiagen). ddPCR was performed using the QX200 Droplet Digital PCR System (Bio-Rad), as previously described^25^ and the *KRAS* G12/G13 and the *BRAF* V600 Screening Multiplex Kits (Bio-Rad). More information on ddPCR is provided in the SI-S4.

## 3. Results and discussion

### Concept of combined AS-PCR / acoustic detection

The main objective of this work was to design a methodology for liquid and tissue biopsy that could exhibit the high sensitivity of ddPCR using a less cumbersome and more cost-effective method. The protocol we developed involves the specific amplification of the point mutation via AS-PCR, followed by acoustic detection using a novel array biochip device. The basic principle of the methodology is presented in Fig. 1A and B.

**Figure 1.**
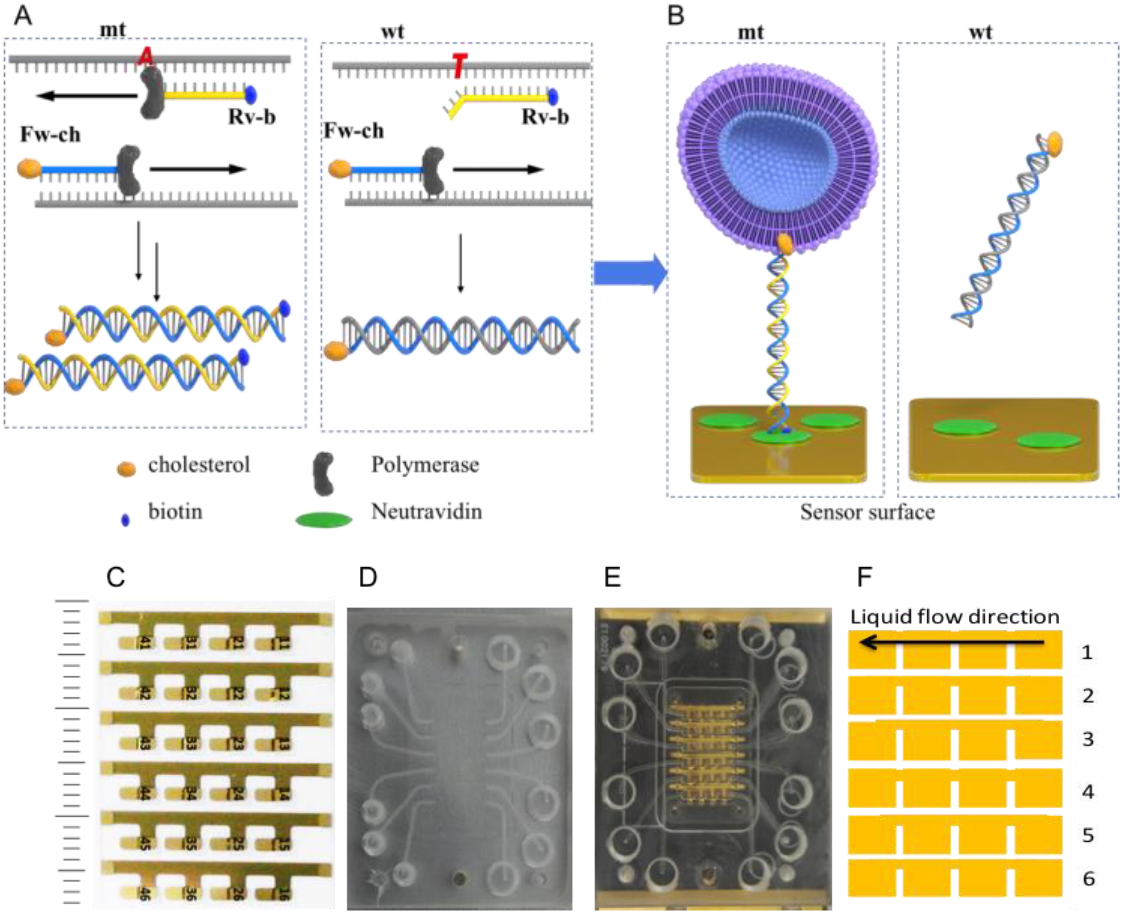
Schematic illustration of the concept of the AS-PCR (A) and acoustic detection (B) for the *BRAF* V600E point-mutation. (C) The acoustic array consists of 24 HFF-QCM sensors arranged in 6 lines of 4 sensors; (D) The PDMS flow cell alone and (E) integrated with the array/PCB board; (F) Schematic representation of the liquid flow along the 6 lines and over the 4 sensors.

For the specific amplification of the mutant allele with AS-PCR, a set of primers^26^ was used of which the forward (Fw) amplifies both the mutant (mt) and wild type (wt) sequences while the reverse (Rv) only the mt target; this is due to the Rv design which has the allele-specific (AS) nucleotide for the mt target in the last position of the 3’-end. To further enhance Rv primer’s specificity, an artificial mismatched nucleotide was introduced at the third from the 3’-end position. For the sake of the downstream acoustic analysis, the Fw primer is modified with a cholesterol in its 5’-end and the Rv primer by a biotin (Fig. 1A).

Following amplification, DNA fragments of 89 bp employing both a biotin and a cholesterol molecule in the case of the mt target or only a cholesterol molecule in the case of the wt target are produced. The AS-PCR reaction is then loaded directly on the NAv-modified acoustic biochip without prior purification. The presence of NAv allows the immobilization of only the mt DNA-amplicons which carry the biotin. In order to achieve the clinically relevant detection limits of few copies of mt target in the presence of larger amounts of the wt (from 0.001-5%), ultrasensitive detection is necessary, even after AS-PCR. This becomes even more significant if detection occurs directly inside the crude AS-PCR cocktail where issues of non-specific binding become of concern. For this reason, a solution of POPC liposomes is injected and captured by the immobilized products via the cholesterol-end of the mt amplified products (Fig. 1B). Liposomes act as signal enhancers causing big changes in the acoustic signal leading to the detection of immobilized DNA. This strategy has been shown by our group as well as others to be suitable for the acoustic detection of Recombinase Polymerase Amplification (RPA) products^27^ and single-base mismatches^28–30^.

### Acoustic array biochip for multiple sample analysis

The HFF resonator array used in this work includes 24 miniaturized crystals integrated monolithically to a single substrate, in a layout of six rows with four resonators/row (Fig. 2C). The HFF QCM array chip is ideal for high-throughput analysis^31^ and low-volume biosensing applications. Because the array is very small and fragile for direct handling during experiments, it was mounted on a PCB (Fig. 2E); the latter also provides mechanical, electrical and thermal interface between the acoustic wave device and the recording instrument. A gasket and a cell have been developed and integrated with the PCB + array assembly (Fig. 2D & E). The flow cell device seals the microsensors individually, so that it is possible to flow liquid in the desired direction over the sensor top surface without affecting the array electrical connections placed on the bottom surface or interfere with the different lines of sensors. During experimental conditions the liquid moves sequentially on each one of the four sensors in the same row (Fig. 2F), running from the input to output. Each crystal has 0.3114 mm^2^ active surface area and 1.5 μL volume above the sensor. With the current flow set-up, six samples can be analyzed in a semi-parallel way with the possibility to perform four tests per sample. More information on the array can be found in Fernandez R.^32^.

**Figure 2.**
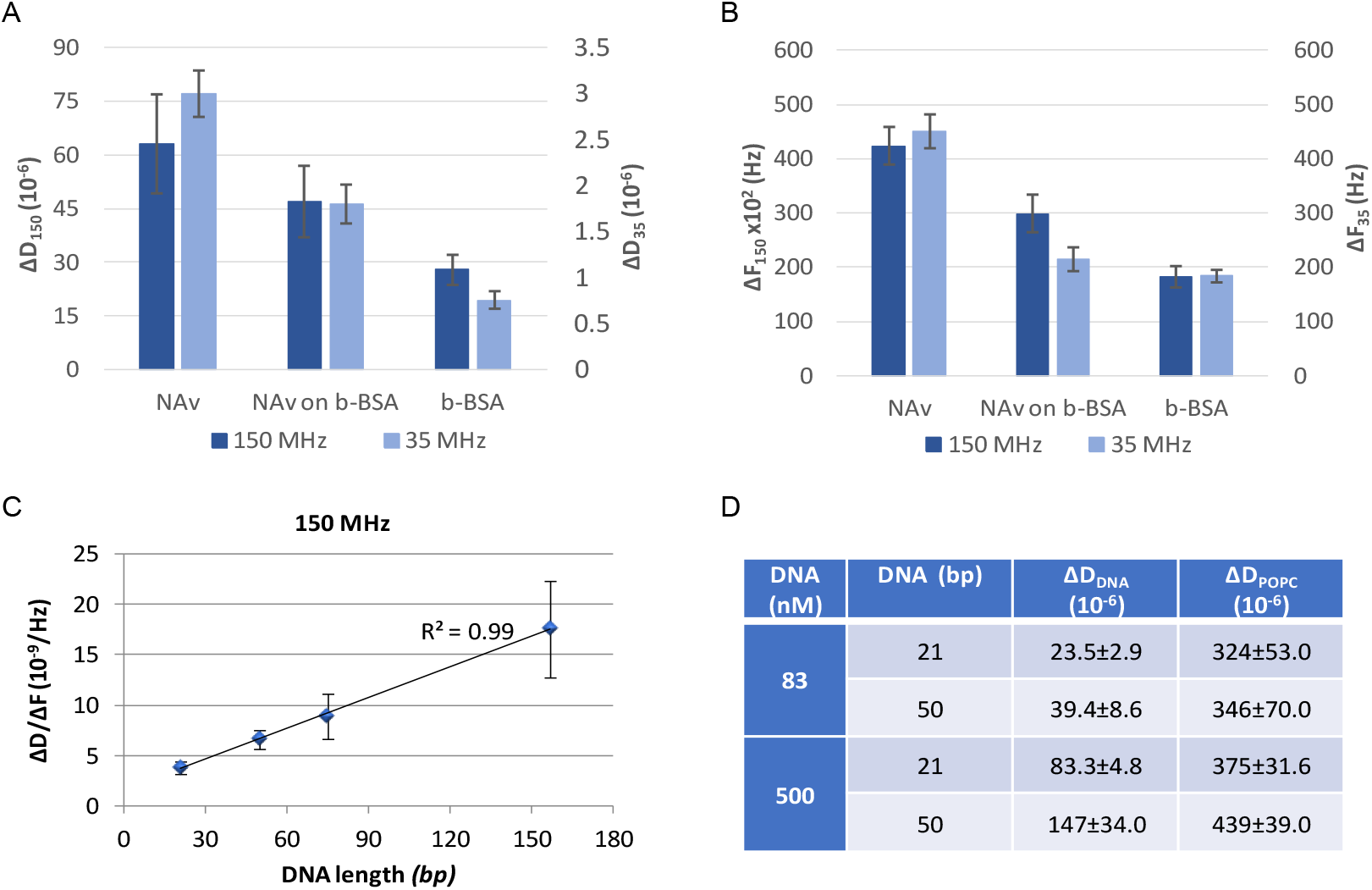
Comparison of ΔD (A) and ΔF values (B) of NAv and b-BSA adsorption as well as NAv binding to pre-adsorbed b-BSA on the 150 MHz and 35 MHz sensors; (C) Acoustic ratio (ΔD/ΔF) as a function of the length of b-DNA attached on a NAv-modified surface; (C) Table of ΔD values at saturation during the binding of DNA (83 & 500 nM) followed by the addition of liposomes (200 nm).

### Performance evaluation of the biochip array

Prior to the development of the combined AS-PCR/acoustic biochip detection assay, we investigated the performance of the device in the presence of pure samples and during the detection of protein, DNA and liposomes-binding. Overall, the principle of operation of the HFF QCM resonator is the same to that of a typical acoustic device; briefly, the presence of an analyte on the sensor surface affects the propagation characteristics of the acoustic wave, i.e. its velocity and energy, which in turn are expressed as changes in frequency (ΔF) and dissipation (ΔD). ΔF changes correlate with the amount of the deposited mass on the sensor^33^; ΔD changes reflect, among other things, the viscoelastic properties of the surface-attached layer and the acoustic ratio (ΔD/ΔF) correlates to the hydrodynamic properties^34–36^ or conformation^37–40^ in the case of discretely-bound molecules. We firstly analyzed the device response during the absorption of proteins as well as the detection of dsDNA and compared them to the response of the standard 35MHz QCM-D. Specifically, the physisorption of NAv and b-BSA protein directly on gold was first recorded as well as the subsequent binding of NAv on a preadsorbed layer of b-BSA. After each addition step, buffer rinsing was performed and the ΔF and ΔD changes were measured at equilibrium and at surface saturation. Figure 2A & 2B shows that the relative signal responses of all three proteins upon absorption/binding to the surface is the same for both devices. Moreover, the array biochip was tested and found to give the expected linear relationship between the DNA length and the acoustic ratio (ΔD/ΔF) during the binding of b-DNA molecules to a NAv-covered surface through a single point (Fig. 2C)^37,39^.

The protocol we developed for the detection of cancer point mutations in crude samples includes a step of amplification through liposomes; for this reason, we further monitored the binding of 200 nm diameter POPC liposomes on dsDNA in buffer. We tested two different lengths of DNA i.e., 21 bp and 50 bp carrying a biotin at their 5’-end for immobilization to the surface and a cholesterol at their 3’-end for the binding of liposomes. ΔD values measured with the 150 MHz QCM array biochip during liposome attachment gave a higher dissipation value when liposomes were bound to the longer DNA (Fig. 2D), in agreement with previous studies^24^. Overall, the successful detection of proteins, DNA and liposomes with the 150 MHz acoustic array demonstrate the suitability of the new system for subsequent application for the development of molecular biology assays in crude samples.

### Analytical performance of the AS-PCR / acoustic assay for the detection of point mutations using genomic DNA

#### (A) BRAF V600E

To determine the limit of detection (LOD) and sensitivity of the assay, mt genomic DNA carrying the *BRAF* V600E point-mutation was mixed with wt DNA in a range from 0.01% to 10% (i.e., from 1:10^4^ to 1:10 mt:wt). The mt:wt dilutions as well as the 100% (10^4^ copies) wt genomic DNA (control) were subjected to 55 cycles of AS-PCR (1h 40 min) followed by acoustic detection on the biochip array. For the immobilization of the biotinylated mt target, we used a surface pre-modified with b-BSA/NAv which was shown in our lab to have a high stability in the presence of a crude sample (data not shown).

Initially, we investigated whether the ΔF and ΔD signals obtained from the direct detection of the immobilization of the AS-PCR products on the b-BSA/NAv-surface could differentiate between the mt and wt reactions. Results indicated poor discrimination between the two (Fig. S1). We attributed this response to the non-specific adsorption of components in the PCR cocktail (e.g., primer dimers, DNA polymerase etc.) resulting in a background signal over-shadowing the response from the specific binding of the biotinylated amplicons. For this reason, we employed a second step, where liposomes were used as signal amplifiers upon binding at the 5’ cholesterol present at the mt amplicon. The real-time binding of the liposomes to the AS-PCR modified surface was specific, giving nearly zero dissipation change in the absence of the mt target (Fig. 3A). Figure 3C summarizes the average dissipation values recorded at the liposome step for all the mt:wt dilutions; based on this data, the sensitivity and LOD for the detection of *BRAF* V600E is 0.01% mt:wt copies and 1 copy, respectively. For the same samples, the frequency response was not as sensitive as the dissipation one, giving large error bars and a lower discrimination capability (Fig. S2). This is in agreement with previous works, where the dissipation signal was proven to be more sensitive to frequency for the ultrasensitive detection of DNA (fmol) via liposomes as signal amplifiers^27,30^.

**Figure 3.**
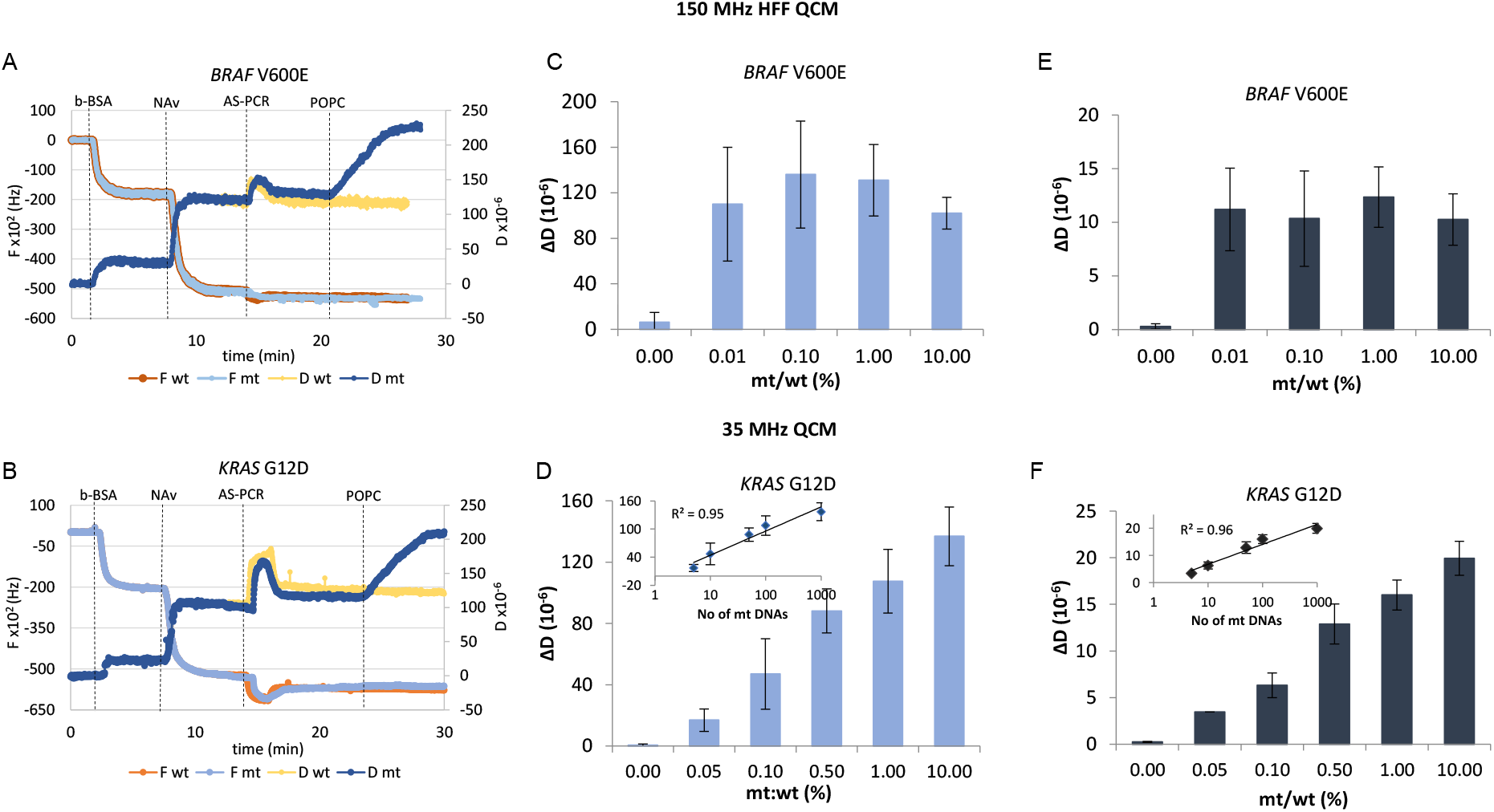
(A) Real time acoustic detection of the *BRAF* V600E mutation together with a control sample (10^4^ wt DNAs) with the 150 MHz acoustic biochip array; (B) As in (A)for the *KRAS* G12D; (C) Recorded acoustic values (saturation) during the detection of the *BRAF* V600E mixed with wt DNA in a range from 0% to 10% using the 150 MHz array; (D) As in (C) for the *KRAS* G12D. The inset shows the linear curve relationship (*R^2^*= 0.95) of the obtained ΔD values when plotted vs the number of mt molecules in logarithmic scale; (E) and (F) as in (C) and (D) for the 35 MHz QCM device. The 0.00% corresponds to the control (10^4^ wt molecules).

Figure 3C shows that the assay provides only qualitative results since it can differentiate between the presence or absence of the mutation but cannot distinguish between the various mt:wt ratios. This is attributed to the fact that the AS-PCR reaction had reached a plateau after the 55 cycles employed in this assay.

#### (B) KRAS G12D

The combined AS-PCR/acoustic methodology was further applied to the detection of the most frequently observed *KRAS* mutation in cancer patients, the G12D, a G→A transition^41^. For this assay, a slightly modified version of Fig. 1A was employed: here, the Rv primer was modified by a cholesterol and designed to amplify both the mt and wt targets, while the Fw was biotinylated and was specific for the mutant allele. Similarly, to the *BRAF*, the analytical performance of the assay was investigated by mixing genomic DNA carrying the *KRAS* G12D point-mutation with wt DNA at gradually decreasing ratios ranging from 0.05% to 10%. The mt:wt dilutions and control of 100% (10^4^ copies) wt genomic DNA were subjected first to real-time AS-PCR; this was in order to determine the number of cycles needed for all the samples to be at the range of the exponential-to-early-plateau phase of the PCR relatively to their initial mt DNA input. Based on these results (Fig. S3), a modified protocol for the AS-PCR including 45 cycles (1h) was established and used for the KRAS detection. For the acoustic analysis, liposome addition on the biochip array pre-loaded with the mt:wt solutions allowed the clear distinction between the mt from the wt samples (Fig. 3B & D). Moreover, the ΔD values obtained upon liposome addition were analogous to the initial mt:wt ratio; the linear relationship obtained between ΔD and the absolute number (logarithmic) of mt molecules (*R^2^*= 0.95) indicates that the method is quantitative (Fig. 3D inset), with a sensitivity and LOD of 0.05% and 5 molecules, respectively. Regarding frequency change, the response was not as sensitive as the dissipation (Fig. S4), similarly to the *BRAF* V600E results.

A notable difference to the *BRAF* analysis was that the ΔD measurement obtained from the direct binding of the AS-PCR products on the b-BSA/NAv coated surface could discriminate to some extent the mt:wt ratios from the wt, although the error bars were very large (Fig S5). This result can be attributed to the higher volume used for the *KRAS* AS-PCR reaction as opposed to the *BRAF* one (20 μl vs 10 μl) and for loading on the HFF QCM sensor surface (8 μl vs 2.5 μl). Moreover, the reduction of the AS-PCR cycles to 45 may have also resulted in fewer by-products and lower non-specific binding. Overall, the above results indicate that upon optimization of the assay conditions, ultra-sensitive, specific and quantitative information could be provided through acoustic measurement of the dissipation signal.

### Effect of the operating frequency

To evaluate the efficiency of the 150 MHz acoustic biochip array towards the analysis of cancer point mutations we used the established 35MHz QCM sensor to perform the same assays and compare results. Differences between the two devices would include the penetration depth inside the sensing solution (43 nm for the 150 MHz and 90 nm for the 35 MHz) and the sensitivity of frequency to the adsorbed mass; the latter according to the Sauerbrey equation scales up with the square of the fundamental resonator frequency fo. In addition, differences in the size of the two QCM-devices, geometry of the flow cell and applied flow rate can affect the amount of immobilized target.

Based on Fig. 3E and F, we conclude that the two devices give the same dissipation response and LOD towards the detection of all tested mt:wt AS-PCR samples. Moreover, the 35 MHz device also failed to detect the *BRAF* and *KRAS* AS-PCR samples upon their direct loading on the b-BSA/NAv coated sensor (Fig. S6 & S7).

### Clinical validation of the method during the analysis of *BRAF* and *KRAS* mutations in tissue samples

The final goal of this work was to assess the capability of the method to detect ctDNA carrying point mutations in patients’ samples. For this reason, initially, we evaluated the ability of the combined AS-PCR/acoustic method to detect *BRAF* V600E and *KRAS* G12D mutant alleles in clinical FFPE tissue samples based on the protocols described before. Regarding the *BRAF* V600E mutation analysis, a total of 11 samples collected from lung (L), melanoma (MEL) and colorectal cancer (CRC) patients’ tissues as well as healthy individuals were tested. Of the above, 6 were positive and 5 negative, as identified by Sanger sequencing and ddPCR. For the KRAS mutation, 10 samples in total were tested, 6 mt and 4 wt, also derived from patients and healthy donors and identified as before. Results are summarized in Table 1 for Sanger sequencing, ddPCR and AS-PCR/acoustic detection. In addition, the calculated mutant allele frequency (MAF) is provided, where MAF is defined as the % of mt/(mt+wt) for each sample, based on ddPCR-measurement of the absolute No of mt and wt copies in each sample.

**Table 1.** Analysis of patients’ FFPE samples with AS-PCR/acoustic methodology, Sanger sequencing and ddPCR. Light and dark grey columns correspond to *BRAF* and *KRAS* samples, respectively. MEL-Melanoma; L-Lung; CRC-Colorectal; MAF-Mutant Allele Frequency.

| Tissue sample | Acoustic $\Delta D$ ($\times 10^{-6}$) | Sanger | ddPCR | | MAF% |
| --- | --- | --- | --- | --- | --- |
|  |  |  | mt copies | wt copies |  |
| **1906 (MEL)** | **44 ± 10** | **mt** | **960** | **914** | **51.2** |
| **1947(L)** | 0 | wt | 0 | 3040 | 0 |
| **2786 (CRC)** | **120 ± 17** | **mt** | **262** | **560** | **31.8** |
| **2840 (L)** | 0 | wt | 0 | 2600 | 0 |
| **4458 (L)** | 0 | wt | 0 | 3100 | 0 |
| **6316 (L)** | 0 | wt | 0 | 4480 | 0 |
| **7874 (L)** | 0 | wt | 0 | 410 | 0 |
| **10014 (MEL)** | **110 ± 31** | **mt** | **448** | **666** | **40.2** |
| **142648 (MEL)** | **138 ± 25** | **mt** | **1994** | **2880** | **40.9** |
| **168078 (L)** | **224 ± 51** | **mt** | **1052** | **1706** | **38.1** |
| **168906 (CRC)** | **47 ± 3.9** | **mt** | **544** | **800** | **40.4** |
| **K534 (CRC)** | 0 | wt | 0 | 3860 | 0 |
| **10434 (CRC)** | 0 | wt | 0 | 1606 | 0 |
| **6229-10 (L)** | **125 ± 9** | **mt** | **834** | **1384** | **37.6** |
| **7102 (CRC)** | **119 ± 11** | **mt** | **222** | **244** | **47.6** |
| **7874 (L)** | 0 | wt | 0 | 592 | 0 |
| **10272 (L)** | 0 | wt | 0 | 1540 | 0 |
| **10732 (CRC)** | **122 ± 15** | **mt** | **640** | **704** | **47.6** |
| **12784 (CRC)** | **135 ± 21** | **mt** | **390** | **592** | **39.7** |
| **164665 (CRC)** | **137 ± 16** | **mt** | **1752** | **1774** | **49.6** |
| **159216 (CRC)** | **104 ± 24** | **mt** | **774** | **1396** | **35.7** |

\*mt, wt copies refer to the number of molecules per 50 ng input DNA.

Based on Table 1, the obtained acoustic results were in full agreement with the Sanger sequencing and ddPCR. Overall, no false positives or false negatives were recorded, indicating 100% sensitivity and specificity for both targets. Moreover, all wt samples gave a zero response, indicating the power of the proposed acoustic methodology to discriminate at least qualitatively (positive/negative result) between malignant and benign tissues.

Finally, the analysis of the samples shown in Table 1 based on the frequency measurement failed to discriminate malignant from healthy tissue samples.

### Clinical validation during *KRAS* and *BRAF* mutations detection in patients’ plasma samples

The method was further evaluated for the detection of ctDNA targets carrying the *BRAF* V600E or *KRAS* G12D mutation using cfDNA derived from patients’ plasma samples. In practice, 10 plasma samples identified as *BRAF* V600E or *BRAF* wt by ddPCR were subjected to AS-PCR followed by acoustic detection; for these experiments, 50 cycles (1 ½ hr) of AS-PCR were used instead of 55, following further optimization of the protocol. In contrast to results obtained with tissue samples, in the case of plasma a change in ΔD was detected for the wt specimens, although significantly lower to that obtained for positive samples. We first tried to identify the source of the background signal; gel electrophoresis of all tested wt samples indicated a high degree of by-products even after 50 PCR cycles. We attributed this response to the presence of cholesterol-primers aggregation, possibly due to the specific batch used in these experiments. To define the cut off value above which ΔD change would be considered as an indication of a positive sample, we calculated the mean ΔD value from all wt samples (healthy) plus three standard deviations^20^; this cut off value was set at a ΔD > 80×10^-6^. Based on the above criterion and results presented in Table 2, we concluded that all samples carrying the *BRAF* V600E point mutation were correctly identified with our method as positive and discriminated from samples coming from healthy individuals. Statistical analysis verified that the mt:wt populations were significantly different (p<0.001). Regarding the *KRAS* G12D mutation analysis, 10 samples identified as *KRAS* G12D or *KRAS* wt by ddPCR were blindly tested by the AS-PCR-acoustic detection, as well. According to Table 2, the combined AS-PCR/acoustic method provided results in full agreement with the ddPCR. Note that, two samples, the 11MEL and 1797L ones, corresponding to 27 and 19 *BRAF* and *KRAS* mutant copies respectively, and a MAF value of <1% were clearly identified as positive by the acoustic method.

**Table 2.** Analysis of patients’ plasma samples with ddPCR and AS-PCR/acoustic detection. Light and dark grey columns correspond to *BRAF* and *KRAS* samples, respectively. MEL-Melanoma; L-Lung; CRC-Colorectal; MAF-Mutant Allele Frequency.

| Plasma sample | Acoustic $\Delta D$ ($\times 10^{-6}$) | ddPCR | | MAF% |
| --- | --- | --- | --- | --- |
|  |  | mt copies | wt copies |  |
| 2 (MEL) | 166 ± 21 | 6110 | 4190 | 59.2 |
| 3 (MEL) | 40 ± 2 | 0 | 194 | 0 |
| 4 (MEL) | 230 ± 35 | 133 | 346 | 27.7 |
| 5 (MEL) | 35 ± 8 | 0 | 389 | 0 |
| 9 (MEL) | 20 | 0 | 7050 | 0 |
| 10 (MEL) | 9 ± 5 | 0 | 523 | 0 |
| 11 (MEL) | 97 ± 13 | 27 | 2680 | 0.99 |
| 14 (MEL) | 55 | 0 | 865 | 0 |
| 1845 (L) | 212 ± 53 | 67 | 1345 | 4.7 |
| 1871 (L) | 199 ± 56 | 100 | 5490 | 1.78 |
| 1504 (CRC) | 123 ± 3 | 1432 | 8940 | 13.8 |
| 1523 (CRC) | 0 | 0 | 163 | 0 |
| 1574 (CRC) | 0 | 0 | 698 | 0 |
| 1596 (CRC) | 118 ± 2.8 | 162 | 1198 | 11.91 |
| 1597 (CRC) | 175 ± 35 | 1574 | 1838 | 46.13 |
| 1602 (CRC) | 0 | 0 | 2500 | 0 |
| 1604 (CRC) | 3 ± 3.5 | 0 | 1594 | 0 |
| 1618 (CRC) | 113 ± 27 | 32.8 | 922 | 3.43 |
| 1797 (L) | 21 ± 2 | 19 | 2080 | 0.91 |
| 1827 (L) | 92 ± 5 | 70.6 | 910 | 7.19 |

\*mt, wt copies refer to the number of molecules per AS-PCR reaction, as calculated by ddPCR.

Finally, as in genomic and FFPE tissue samples analysis, frequency response of the AS-PCR followed by acoustic detection at 150 MHz was not as sensitive as the dissipation (Fig. S8).

### Comparison of the combined AS-PCR / acoustic method to current state-of-the-art for liquid biopsy

Allele specific PCR has been used extensively for the detection of mutations in serum and plasma. So far, both the traditional assay and its variants, e.g., the AS-NEPB-PCR^42^, cAS-PCR^26^, cAS-PCR (rcAS-PCR), CAST-PCR)^43^ and ARMS^44–46^, have limited analytical sensitivity in the range of 0.1-2%, affecting their potential broad clinical use^47^. In all the above assays, detection takes place in situ, i.e. during amplification, using fluorescent probes. Compared to the above performance, the acoustic detection of the AS-PCR amplicons seems to be a promising alternative reaching a detection capability of 0.01% mutant alleles without increasing the complexity of the assay. We attribute the enhanced sensitivity of the acoustic methodology in the two-step assay employed, i.e., first the direct immobilization of the amplicons on the device surface followed by acoustic signal enhancement (here liposomes). Desirable features of the proposed method are the use of double-labeled primers allowing the direct binding of double stranded DNA on the device, by-passing the need for denaturation and surface hybridization as well as the high sensitivity of the dissipation signal which can detect fmol of target DNA^24^.

Currently one of the gold standards in a clinical oncology lab is the use of the real-time PCR based COBAS mutation test; the method designed for tissue (KRAS, EGFR mutations) and liquid (EGFR) biopsy allows simultaneous detection of FFPE tissues and plasma samples within <8 and 4 h respectively and detects ≥ 5% mutant allele copies in a background of wild-type DNA. Our proposed technique outperforms the above commercially available method in terms of both sensitivity (0.01-0.05%) and total analysis time (<5 h and <3 h for tissue and plasma, respectively). However, the COBAS test offers the advantages of multiple samples analysis and ability to detect a panel of mutations (19 KRAS, 42 EGFR) simultaneously. With the current acoustic biochip design, six samples can be detected per array (giving 4 readings per sample) as demonstrated during the detection of two point mutations. Employing a new flow-cell design with the current array, to test 8 (3 readings/sample) or 16 (2 readings/sample) mutations can improve multiple analysis. In addition, given the low cost of the biochip array (<1$ for large scale production), 2 or 3 array biochips could be used in parallel, without increasing substantially the complexity and size of the instrumentation.

Comparing to the ddPCR, our method is more cost-effective since we use a standard thermocycler (<2.6K) and the acoustic platform (<10K), both an order of magnitude less expensive than a ddPCR machine (110K). The development of a single platform integrating a thermocycling unit with acoustic detection is also a feasible future plan.

## 4. Conclusions

In this work, we report the successful application of a novel HFF-QCM 24-sensors array biochip for the acoustic detection of *BRAF*-V600E and *KRAS*-G12D point-mutations from patients’ tissue and plasma samples upon enzymatic amplification. AS-PCR was chosen for amplification due to the method’s high specificity and already wide applicability on a routine basis. The excellent sensitivity of the acoustic method (0.01-0.05% MAF), shown here to be comparable to that obtained with ddPCR but for a fraction of the cost and in a much faster manner, together with its technology-readiness level hold promise for fast adaptation in a clinical oncology lab for both tissue and liquid biopsy. It is anticipated that the proposed methodology can become a promising tool to identify patients carrying somatic mutations in *BRAF* or *KRAS* genes.

## Acknowledgements

This work was supported by the European Union’s Horizon H2020-FETOPEN-1-2016-2017 under grant agreement No. 737212 (CATCH-U-DNA).

## Appendix A. Supplementary material

## REFERENCES

1. Heitzer, E., Haque, I. S., Roberts, C. E. S. & Speicher, M. R. Current and future perspectives of liquid biopsies in genomics-driven oncology. Nat. Rev. Genet. 20, 71–88 (2019).

2. Han, X., Wang, J. & Sun, Y. Circulating Tumor DNA as Biomarkers for Cancer Detection. Genomics, Proteomics and Bioinformatics vol. 15 59–72 (2017).

3. Parikh, A. R. et al. Liquid versus tissue biopsy for detecting acquired resistance and tumor heterogeneity in gastrointestinal cancers. Nat. Med. 25, 1415–1421 (2019).

4. Soda, N., Rehm, B. H. A., Sonar, P., Nguyen, N. T. & Shiddiky, M. J. A. Advanced liquid biopsy technologies for circulating biomarker detection. J. Mater. Chem. B 7, 6670–6704 (2019).

5. Crowley, E., Di Nicolantonio, F., Loupakis, F. & Bardelli, A. Liquid biopsy: Monitoring cancer-genetics in the blood. Nat. Rev. Clin. Oncol. 10, 472–484 (2013).

6. Elazezy, M. & Joosse, S. A. Techniques of using circulating tumor DNA as a liquid biopsy component in cancer management. Computational and Structural Biotechnology Journal vol. 16 370–378 (2018).

7. Gorgannezhad, L., Umer, M., Islam, M. N., Nguyen, N. T. & Shiddiky, M. J. A. Circulating tumor DNA and liquid biopsy: Opportunities, challenges, and recent advances in detection technologies. Lab on a Chip vol. 18 1174–1196 (2018).

8. Acinas, S. G., Sarma-Rupavtarm, R., Klepac-Ceraj, V. & Polz, M. F. PCR-induced sequence artifacts and bias: Insights from comparison of two 16s rRNA clone libraries constructed from the same sample. Appl. Environ. Microbiol. 71, 8966–8969 (2005).

9. Kanagawa, T. Bias and artifacts in multitemplate polymerase chain reactions (PCR). J. Biosci. Bioeng. 96, 317–323 (2003).

10. Wang, Q. et al. Electrochemical biosensors for detection of point mutation based on surface ligation reaction and oligonucleotides modified gold nanoparticles. Anal. Chim. Acta 688, 163–167 (2011).

11. Nur, S., Kosova, B. & Ozsoz, M. Detection of Janus Kinase 2 gene single point mutation in real samples with electrochemical DNA biosensor. Clin. Chim. Acta 429, 134–139 (2014).

12. Zeng, N. & Xiang, J. Detection of KRAS G12D point mutation level by anchor-like DNA electrochemical biosensor. Talanta 198, 111–117 (2019).

13. Sierpe, R., Kogan, M. J. & Bollo, S. Label-free oligonucleotide-based spr biosensor for the detection of the gene mutation causing prothrombin-related thrombophilia. Sensors (Switzerland) 20, 1–13 (2020).

14. Nguyen, A. H. & Sim, S. J. Nanoplasmonic biosensor: Detection and amplification of dual bio-signatures of circulating tumor DNA. Biosens. Bioelectron. 67, 443–449 (2015).

15. Papadakis, G. & Gizeli, E. Screening for mutations in BRCA1 and BRCA2 genes by measuring the acoustic ratio with QCM. Anal. Methods 6, 363–371 (2014).

16. Papadakis, G. et al. Acoustic detection of DNA conformation in genetic assays combined with PCR. Sci. Rep. 3, 1–8 (2013).

17. Dell’Atti, D., Tombelli, S., Minunni, M. & Mascini, M. Detection of clinically relevant point mutations by a novel piezoelectric biosensor. in Biosensors and Bioelectronics vol. 21 1876–1879 (2006).

18. Xu, Q. et al. Detection of single-nucleotide polymorphisms with novel leaky surface acoustic wave biosensors, DNA ligation and enzymatic signal amplification. Biosens. Bioelectron. 33, 274–278 (2012).

19. Bellassai, N. & Spoto, G. Biosensors for liquid biopsy: circulating nucleic acids to diagnose and treat cancer. Analytical and Bioanalytical Chemistry vol. 408 7255–7264 (2016).

20. Das, J., Ivanov, I., Sargent, E. H. & Kelley, S. O. DNA Clutch Probes for Circulating Tumor DNA Analysis. J. Am. Chem. Soc. 138, 11009–11016 (2016).

21. D’Agata, R. et al. Direct plasmonic detection of circulating RAS mutated DNA in colorectal cancer patients. Biosens. Bioelectron. 170, (2020).

22. Kirimli, C., Lin, S., Su, Y., Shih, W. & Shih, W. Y. In situ, amplification-free double-stranded mutation detection at 60 copies / ml with thousand-fold wild type in urine. Biosens. Bioelectron. 119, 221–229 (2018).

23. Alix-Panabieres, C. The future of liquid biopsy. Nature 579, S9 (2020).

24. Milioni, D. et al. Acoustic Methodology for Selecting Highly Dissipative Probes for Ultrasensitive DNA Detection. Anal. Chem. 92, 8186–8193 (2020).

25. Michaelidou, K. et al. Detection of KRAS G12/G13 Mutations in Cell Free-DNA by Droplet Digital PCR, Offers Prognostic Information for Patients with Advanced Non-Small Cell Lung Cancer. Cells 9, 1–18 (2020).

26. Yang, Z. et al. Improved detection of BRAF V600E using allele-specific PCR coupled with external and internal controllers. Sci. Rep. 7, 1–12 (2017).

27. Grammoustianou, A., Papadakis, G. & Gizeli, E. Solid-Phase Isothermal DNA Amplification and Detection on Quartz Crystal Microbalance Using Liposomes and Dissipation Monitoring. IEEE Sensors Lett. 1, 1–4 (2017).

28. Patolsky, F., Lichtenstein, A., Willner, I. & October, R. V. Electronic Transduction of DNA Sensing Processes on Surfaces: Amplification of DNA Detection and Analysis of Single-Base Mismatches by Tagged Liposomes. Am. Chem. Soc. 123, 5194–5205 (2001).

29. Patolsky, F., Lichtenstein, A., Willner, I. & August, R. V. Amplified Microgravimetric Quartz-Crystal-Microbalance Assay of DNA Using Oligonucleotide-Functionalized Liposomes or Biotinylated Liposomes. 418–419 (2000) doi:10.1021/ja992834r.

30. Milioni, D. et al. Acoustic Methodology for Selecting Highly Dissipative Probes for Ultrasensitive DNA Detection. Anal. Chem. 92, 8186–8193 (2020).

31. Mitsakakis, K. & Gizeli, E. Multi-sample acoustic biosensing microsystem for protein interaction analysis. Biosens. Bioelectron. 26, 4579–4584 (2011).

32. Fernandez, R. et al. High Fundamental Frequency (HFF) Monolithic Resonator Arrays for Biosensing Applications: Design, Simulations, and Experimental Characterization. IEEE Sens. J. 21, 284–295 (2021).

33. Mitsakakis, K., Tsortos, A. & Gizeli, E. Quantitative determination of protein molecular weight with an acoustic sensor; significance of specific versus non-specific binding. Analyst 139, 3918–3925 (2014).

34. Reviakine, I., Johannsmann, D. & Richter, R. P. Hearing What You Cannot See and Visualizing What You Hear: (2011) doi:10.1021/ac201778h.

35. Raptis, V., Tsortos, A. & Gizeli, E. Theoretical Aspects of a Discrete-Binding Approach in Quartz-Crystal Microbalance Acoustic Biosensing. Phys. Rev. Appl. 11, 1 (2019).

36. Vázquez-quesada, A. et al. Hydrodynamics of Quartz-Crystal-Microbalance DNA Sensors Based on Liposome Amplifiers. Phys. Rev. Appl. 10, 1 (2020).

37. Tsortos, A., Papadakis, G., Mitsakakis, K., Melzak, K. A. & Gizeli, E. Quantitative determination of size and shape of surface-bound DNA using an acoustic wave sensor. Biophys. J. 94, 2706–2715 (2008).

38. Papadakis, G., Tsortos, A. & Gizeli, E. Acoustic characterization of nanoswitch structures: Application to the DNA holliday junction. Nano Lett. 10, 5093–5097 (2010).

39. Tsortos, A., Papadakis, G. & Gizeli, E. On the Hydrodynamic Nature of DNA Acoustic Sensing. Anal. Chem. 88, 6472–6478 (2016).

40. Tsortos, Achilleas; Grammoustianou, Aristea; Lymbouridou, Rena; Papadakis, George; Gizeli, E. Detection of multiple DNA targets with a single probe using a conformation-sensitive acoustic sensor. Chem. Commun. (2015) doi:10.1039/c0xx00000x.

41. Prior, I. A., Lewis, P. D. & Mattos, C. A comprehensive survey of ras mutations in cancer. Cancer Res. 72, 2457–2467 (2012).

42. Wang, H. et al. Allele-specific, non-extendable primer blocker PCR (AS-NEPB-PCR) for DNA mutation detection in cancer. J. Mol. Diagnostics 15, 62–69 (2013).

43. Barbano, R. et al. Competitive allele-specific TaqMan PCR (Cast-PCR) is a sensitive, specific and fast method for BRAF V600 mutation detection in Melanoma patients. 1–11 (2015) doi:10.1038/srep18592.

44. Lade-Keller, J. et al. Evaluation of BRAF mutation testing methodologies in formalin-fixed, paraffin-embedded cutaneous melanomas. J. Mol. Diagnostics 15, 70–80 (2013).

45. Lang, A. H. et al. Optimized allele-specific real-time PCR assays for the detection of common mutations in KRAS and BRAF. J. Mol. Diagnostics 13, 23–28 (2011).

46. Aung, K. L. et al. Analytical validation of BRAF mutation testing from circulating free DNA using the amplification refractory mutation testing system. J. Mol. Diagnostics 16, 343–349 (2014).

47. Siravegna, G., Marsoni, S., Siena, S. & Bardelli, A. Integrating liquid biopsies into the management of cancer. Nature Reviews Clinical Oncology vol. 14 531–548 (2017).

