## Supplementary material for "Acoustic array biochip combined with allele-specific PCR for multiple cancer mutation analysis in tissue and liquid biopsy"

**Section 1: Materials and Methods**

S1. Acoustic biochip array and QCM sensor. AWS X24 platform (AWSensors S. L., Spain) was used to characterize simultaneously the acoustic response of up to 24 HFF-QCMD sensors integrated in a monolithic array. This QCMD instrument is based on the fixed-frequency phase-shift measurement technique described elsewhere (Fernandez et al., 2021). AWS X24 incorporates several operation modes; here we used a characterization method based on classical impedance spectroscopy that provides both frequency and dissipation information. AWSuite software package (AWSensors S. L., Spain) was used to control the instrument and to register and process the acquired data. Cleaning of the 150 MHz HFF-QCM acoustic array (PCB + array) included treatment with Hellmanex 2%, rinsing with mili-Q water and drying under N_2_ followed by 30 min UV/Ozone (Ossila Ltd, Sheffield) cleaning and EtOH rinse. The flow cell employed a PMMA gasket and a PDMS cell (AWS, S.L. Paterna, Spain), both cleaned with Hellmanex 2 % and rinsed with mili-Q water prior to use. The QCM sensors (AWS, S.L. Paterna, Spain), with a fundamental frequency of 5 MHz, were monitored using the Q-Sense E4 instrument (Biolin Scientific, Sweden) at the 7th overtone (35MHz). The QCM devices were cleaned following the same protocol as described above. Acoustic experiments were carried out under a continuous flow of 20 μL/min for the AWS platform and of 50 μL/min for the Q-Sense instrument, unless otherwise stated.

S2. Acoustic detection of b-BSA and Nav. b-BSA was prepared after incubation of BSA lyophilized powder (Sigma-Aldrich) with biotin-(AC_5_)_2_-Sulfo-Osu linker (Dojindo) in a molar ratio of 1:10 for 1 h and 30 min at RT and purified using the Microcon-30kDa Centrifugal Filter Unit with Ultracel-30 membrane (Merck).

S3. *BRAF* V600E and *KRAS* G12D AS-PCR. Primers for the *BRAF* V600E mutation detection were obtained from Z. Yang et al.(Yang et al., 2017). As template, 50 ng genomic DNA (gDNA) gDNA BRAF-V600E 50% serially diluted with BRAF wt reference standard (Horizon Discovery Ltd) in ratios ranging from 0.01% to 10% was used. For the evaluation of the assay in clinical samples, 50 ng gDNA extracted from FFPE tissue samples were added. Regarding liquid biopsy, 2 - 3.5 μL extracted cell-free DNA were used as template in the presence of 2 ng background wt genomic DNA. The cycling protocol of the BRAF *V600E* AS-PCR consisted of 50 cycles for cell-free DNA or 55 cycles for gDNA, both 10 sec at 94 ^o^C, 10 sec at 60 ^o^C and 10 sec at 72 ^o^C. A no-template control (NTC) was included in every run. Regrading *KRAS* G12D, as template we used 50 ng gDNA KRAS-G12D 50% serially diluted with KRAS wt reference standard (Horizon Discovery Ltd) in ratios ranging from 0.05% to 10%. Similarly, for clinical sample testing, 50 ng gDNA extracted from FFPE tissue samples were added to the AS-PCR mix. For liquid biopsy 3.5 μL extracted cfDNA were used. PCR reaction mixtures were carried out at CFX Real-Time PCR Detection Systems (Bio-Rad). The cycling protocol consisted of 45 cycles of 10 sec at 94 ^o^C, 10 sec at 62 ^o^C and 2 sec at 72 ^o^C). A no template control reaction (NTC) was included in every run.

S4. Sanger Sequencing and ddPCR for *BRAF* and *KRAS* mutation analysis. For the ddPCR we used 50 ng of gDNA from FFPE tissue samples or 3.5 μL - 4 μL of cell-free DNA extracted from plasma samples. Every ddPCR run included negative control templates (NCT) and positive controls (i.e., mutant homozygous and heterozygous cancer cell lines, as well as wt samples for *KRAS* G12/13 and *BRAF* V600). The QuantaSoft Analysis Pro Software (Version 1.0.596) was used to assign positive/negative droplets, to calculate target copies/μL of reaction and the mutant allele frequency (MAF) for each sample. A threshold was manually set for each sample, based on positive control samples (mutant and wt for each channel-FAM and HEX, respectively). Wells with less than 10^4^ accepted droplets were excluded from further analyses. A minimum of three positive droplets was used to call a sample positive for the *BRAF* V600E mutation, which is the minimum acceptable value for the ddPCR Poisson precision calculation.

**Table S1:** *BRAF* and *KRAS* primer sequences used for AS-PCR and Sanger sequencing.

| **Primer** | **Sequence (5’🡪 3’)** | **Product size (bp)** |
| --- | --- | --- |
| *BRAF* V600EF (Metabion) | chol-ACTACACCTCAGATATATTTCTTCATG | 89 |
| *BRAF* V600ER (Metabion) | biotin–CCCACTCCATCGAGATTGCT |  |
| *KRAS* G12DF (Metabion) | biotin-CTTGTGGTAGTTGAAGCGGA | 85 |
| *KRAS* G12DR (Metabion) | chol-CATATTCGTCCACAAAATGATTCTG |  |
| *KRAS*_Ex2F (Invitrogen) | TAAGGCCTGCTGAAAATGAC | 165 |
| *KRAS*_Ex2R (Invitrogen) | GTCCTGCACCAGTAATATGC |  |
| *BRAF*_Ex15F (Invitrogen) | TGTTTTCCTTTACTTACTACACCTCA | 163 |
| *BRAF*_Ex15R (Invitrogen) | GCCTCAATTCTTACCATCCA |  |

**Section 2: Results**

| mt/wt (%) | ΔD (10-6) |
| --- | --- |
| 0 | 6.65 |
| 0.01 | 10.21 |
| 0.1 | 4.96714 |
| 1 | 10.5987 |
| 10 | 7.63125 |

| mt/wt (%) | ΔF (Hz) |
| --- | --- |
| 0 | 1199.8 |
| 0.01 | 1753.17 |
| 0.1 | 1852.43 |
| 1 | 1774 |
| 10 | 1627.89 |

**Figure S1.** ΔD & ΔF values obtained from the addition of various mt:wt ratios of *BRAF* V600E AS-PCR reactions on b-BSA/NAv coated surface (at 150 MHz). 0.00 % corresponds to AS-PCR containing 0 mt and 10^4^ wt molecules (control).

| mt/wt (%) | ΔF (Hz) |
| --- | --- |
| 0 | -220.6 |
| 0.01 | 1166 |
| 0.1 | 4554.58 |
| 1 | 4246.29 |
| 10 | 1137.33 |

**Figure S2:** Comparison of frequency changes observed after loading of *BRAF* V600E AS-PCR (0.01%, 0.1%, 1%, 10% and 0% mt:wt ratios) on b-BSA/NAv modified sensors followed by the addition of 200 nm diameter POPC liposomes (at 150 MHz). 0.00 % corresponds to AS-PCR containing 0 mt and 10^4^ wt molecules (control).

KRAS-G12D

Flouresence (RFU)

| Cycles | NTC | Wild Type | Mutant 10% | Mutant 1% | Mutant 0,5% | Mutant 0,1% | Mutant 0,05% | Threshold |
| --- | --- | --- | --- | --- | --- | --- | --- | --- |
| 1 | 23.8149 | 126.631 | 216.147 | 184.446 | 155.539 | 549.44 | 280.65 | 1000 |
| 2 | 12.9859 | 74.1075 | 104.279 | 72.2481 | 73.1778 | 387.286 | 182.652 | 1000 |
| 3 | 9.42739 | 77.8496 | 56.8456 | 30.9034 | 54.3765 | 317.98 | 141.986 | 1000 |
| 4 | 6.972 | 57.6016 | 34.3703 | 14.7651 | 36.1834 | 265.645 | 106.449 | 1000 |
| 5 | 4.69333 | 41.5791 | 15.198 | -3.2315 | 19.1738 | 237.283 | 84.2098 | 1000 |
| 6 | 0.708938 | 22.6082 | -12.1067 | -11.8416 | 5.38333 | 188.216 | 65.1917 | 1000 |
| 7 | 0.974431 | 10.779 | -25.7587 | -20.7484 | -4.98472 | 160.145 | 54.8067 | 1000 |
| 8 | -0.480834 | -2.94458 | -38.6961 | -32.2495 | -17.597 | 138.936 | 34.4088 | 1000 |
| 9 | -1.92201 | -10.7663 | -33.653 | -25.7379 | -18.2521 | 111.083 | 21.39 | 1000 |
| 10 | -1.52095 | -22.4777 | -53.4505 | -35.0566 | -28.7671 | 83.4259 | 9.44023 | 1000 |
| 11 | -0.250721 | -21.2523 | -57.9892 | -35.754 | -28.5032 | 52.3137 | -3.16595 | 1000 |
| 12 | -1.89917 | -25.7761 | -65.5421 | -25.5029 | -25.6395 | 34.5946 | -13.4 | 1000 |
| 13 | -1.72349 | -37.2665 | -57.7905 | -24.9472 | -31.1069 | 20.4131 | -24.41 | 1000 |
| 14 | -3.06473 | -42.9963 | -65.5202 | -38.6659 | -40.8311 | -4.29735 | -31.7161 | 1000 |
| 15 | -3.18847 | -49.0798 | -70.9395 | -45.4964 | -47.2881 | -26.7063 | -46.3081 | 1000 |
| 16 | -0.889911 | -59.4573 | -78.2273 | -36.3436 | -47.9005 | -58.8975 | -61.8114 | 1000 |
| 17 | -1.55756 | -62.946 | -72.3063 | -47.3039 | -55.1249 | -73.1715 | -68.829 | 1000 |
| 18 | -3.63191 | -58.2434 | -72.26 | -43.6663 | -50.9548 | -92.2535 | -75.8479 | 1000 |
| 19 | -1.12629 | -67.2041 | -73.215 | -53.9375 | -60.5708 | -111.809 | -79.3709 | 1000 |
| 20 | -0.542009 | -64.2441 | -73.1293 | -48.3846 | -56.3144 | -138.051 | -81.47 | 1000 |
| 21 | -2.365 | -73.568 | -69.9103 | -41.752 | -57.66 | -155.73 | -95.993 | 1000 |
| 22 | -6.81468 | -74.3045 | -59.892 | -42.9357 | -58.6201 | -166.51 | -92.5443 | 1000 |
| 23 | -3.27631 | -78.0945 | -64.7401 | -48.6196 | -63.3571 | -184.91 | -102.589 | 1000 |
| 24 | -1.11005 | -73.2802 | -51.5875 | -42.5899 | -57.9351 | -195.374 | -101.909 | 1000 |
| 25 | -2.84189 | -76.8779 | -50.8578 | -35.9767 | -56.4273 | -216.69 | -99.603 | 1000 |
| 26 | 0.396315 | -74.0781 | -48.6895 | -30.5848 | -52.3315 | -229.199 | -102.608 | 1000 |
| 27 | -1.56589 | -72.7083 | -35.0411 | -31.275 | -51.9916 | -248.21 | -108.179 | 1000 |
| 28 | -2.40754 | -70.0922 | -24.7164 | -29.6704 | -49.8813 | -252.932 | -108.803 | 1000 |
| 29 | 0.198553 | -57.133 | -15.8503 | -19.7095 | -38.4212 | -262.46 | -104.414 | 1000 |
| 30 | -1.69906 | -58.5389 | 6.96915 | -17.5975 | -38.0682 | -273.613 | -109.35 | 1000 |
| 31 | -1.14052 | -59.2972 | 32.7475 | -12.3077 | -35.8025 | -287.29 | -108.55 | 1000 |
| 32 | -2.08509 | -50.0762 | 71.1386 | -8.8768 | -29.4765 | -297.865 | -108.498 | 1000 |
| 33 | 0.164217 | -49.2466 | 125.772 | 3.51963 | -22.8635 | -308.874 | -102.266 | 1000 |
| 34 | 0.0373461 | -48.5699 | 227.457 | 28.1853 | -10.1923 | -310.673 | -98.967 | 1000 |
| 35 | -0.390291 | -36.879 | 413.192 | 66.3397 | 14.7304 | -321.256 | -96.2941 | 1000 |
| 36 | -2.107 | -24.8547 | 720.41 | 125.117 | 50.1311 | -323.382 | -87.1349 | 1000 |
| 37 | -1.12134 | -23.6325 | 1241.63 | 219.937 | 98.1525 | -325.231 | -83.3907 | 1000 |
| 38 | 2.25912 | -15.7819 | 2089.18 | 391.61 | 187.914 | -311.568 | -74.7302 | 1000 |
| 39 | 0.8986 | -2.07921 | 3367.33 | 705.594 | 351.758 | -283.673 | -56.2432 | 1000 |
| 40 | 1.98338 | 8.10293 | 4954.79 | 1244.07 | 626.087 | -224.37 | -31.7265 | 1000 |
| 41 | 1.35591 | 17.5685 | 6486.13 | 2144.25 | 1080.91 | -127.84 | -2.39931 | 1000 |
| 42 | 1.40534 | 23.6849 | 7829.78 | 3553.47 | 1788.58 | 58.4517 | 37.8902 | 1000 |
| 43 | 3.60427 | 36.0279 | 8907.61 | 5463.7 | 2749.87 | 389.57 | 113.278 | 1000 |
| 44 | 2.15334 | 51.0092 | 9774.94 | 7456.36 | 3753.68 | 954 | 243.332 | 1000 |
| 45 | 3.71162 | 60.923 | 10503.8 | 9378.64 | 4719.78 | 1867.82 | 456.768 | 1000 |

**Figure S3:** Schematic representation of the PCR curves obtained by Real-Time AS-PCR of the *KRAS* G12D mt:wt dilutions (0.05%, 0.1%, 0.5%, 1%, 10% and 0% mt:wt ratios). The threshold was set at 1000 RFU (dashed line). In all cases the Real Time PCR assays included a wt reference control which contained 10^4^ wt DNAs (red line) as well as a NTC reaction (blue).

| mt/wt (%) | ΔF (Hz) |
| --- | --- |
| 0 | -341.5 |
| 0.05 | -36.1818 |
| 0.1 | -89.8889 |
| 0.5 | 269.667 |
| 1 | 724.625 |
| 10 | 1693.12 |

**Figure S4:** Comparison of frequency changes observed after loading of the *KRAS* G12D AS-PCR (0.05%, 0.1%, 0.5%, 1%, 10% and 0% mt:wt ratios) on b-BSA/NAv modified sensors followed by the addition of 200 nm diameter POPC liposomes (at 150 MHz). 0.00 % corresponds to AS-PCR containing 0 mt and 10^4^ wt molecules (control).

| mt/wt (%) | ΔD (10-6) |
| --- | --- |
| 0 | 16.0587 |
| 0.05 | 9.9125 |
| 0.1 | 19.8167 |
| 0.5 | 23.0822 |
| 1 | 34.0144 |
| 10 | 60.7563 |

| mt/wt (%) | ΔF (Hz) |
| --- | --- |
| 0 | 5131.53 |
| 0.05 | 5063.38 |
| 0.1 | 3588.44 |
| 0.5 | 4529 |
| 1 | 3523.22 |
| 10 | 6793.5 |

**Figure S5.** ΔD & ΔF values obtained from the addition of various mt:wt ratios of *KRAS* G12D AS-PCR reactions on b-BSA/NAv coated surface at 150 MHz. 0.00 % corresponds to AS-PCR containing 0 mt and 10^4^ wt molecules (control).

| mt/wt (%) | ΔD (10-6) |
| --- | --- |
| 0 | 0.265 |
| 0.01 | 0.146667 |
| 0.1 | 0.216667 |
| 1 | 0.116667 |
| 10 | 0.132333 |

| mt/wt (%) | ΔF (Hz) |
| --- | --- |
| 0 | 2.89 |
| 0.01 | 1.78 |
| 0.1 | 4 |
| 1 | 1.08333 |
| 10 | 2.7689 |

**Figure S6.** ΔD & ΔF values obtained from the direct loading of various mt:wt ratios of *BRAF* V600E AS-PCR reactions on b-BSA/NAv coated surface (at 35 MHz). 0.00 % corresponds to AS-PCR containing 0 mt and 10^4^ wt molecules (control).

| mt/wt (%) | ΔD (10-6) |
| --- | --- |
| 0 | 0.1025 |
| 0.05 | 0.1736 |
| 0.1 | 0.12725 |
| 0.5 | 0.315 |
| 1 | 0.4575 |
| 10 | 0.52 |

| mt:wt (%) | ΔF (Hz) |
| --- | --- |
| 0 | 5.54111 |
| 0.05 | 3.5 |
| 0.1 | 4.4275 |
| 0.5 | 9.4275 |
| 1 | 9.625 |
| 10 | 14.7 |

**Figure S7.** ΔD & ΔF values obtained from the direct loading of various mt:wt ratios of *KRAS* G12D AS-PCR reactions on b-BSA/NAv coated surface (at 35 MHz). 0.00 % corresponds to AS-PCR containing 0 mt and 10^4^ wt molecules (control).

**B**

**A**

| Sample | ΔF (Hz) |
| --- | --- |
| 3MEL | -464 |
| 5MEL | -2807 |
| 9MEL | -1361 |
| 10MEL | 473 |
| 14MEL | 350 |
| 2MEL | 1317 |
| 4MEL | 1871 |
| 11MEL | 345 |
| 1871L | -557 |
| 1845L | 1662 |

| Sample | ΔF (Hz) |
| --- | --- |
| 1523CRC | -82.33 |
| 1604CRC | -163 |
| 1574CRC | 355.7 |
| 1602CRC | -140 |
| 1797L | 716 |
| 1618CRC | 4646 |
| 1827L | 1810 |
| 1596CRC | 2461 |
| 1504CRC | 1734 |
| 1597CRC | 3581 |

**Figure S8:** Comparison of ΔF values obtained from the analysis of patients’ (A) *BRAF* V600E and (B) *KRAS* G12D plasma samples by AS-PCR & acoustic detection (at 150 MHz). Blue and red columns represent wt and mt samples, respectively.

**References**

Fernandez, R., Calero, M., Garcia-Narbon, J.V., Reiviakine, I., Arnau, A., Jimenez, Y., 2021. A Fast Method for Monitoring the Shifts in Resonance Frequency and Dissipation of the QCM Sensors of a Monolithic Array in Biosensing Applications. IEEE Sens. J. 21, 6643–6651. https://doi.org/10.1109/JSEN.2020.3042653

Yang, Z., Zhao, N., Chen, D., Wei, K., Su, N., Huang, J.F., Xu, H.Q., Duan, G.J., Fu, W.L., Huang, Q., 2017. Improved detection of BRAF V600E using allele-specific PCR coupled with external and internal controllers. Sci. Rep. 7, 1–12. https://doi.org/10.1038/s41598-017-14140-2
